# Identification of DNA response elements regulating expression of CCAAT/enhancer-binding protein (C/EBP) β and δ during early adipogenesis

**DOI:** 10.1101/2020.05.22.110114

**Authors:** James E. Merrett, Tao Bo, Peter J. Psaltis, Christopher G. Proud

## Abstract

Given the high and increasing prevalence of obesity and associated disorders, such as type-2 diabetes, it is important to understand the mechanisms that regulate lipid storage and the differentiation of fat cells, a process termed adipogenesis. Using the well-established mouse 3T3-L1 *in vitro* model of adipogenesis, we refine how the induction of two key adipogenic transcription factors, CCAAT/enhancer-binding proteins (C/EBPs) β and δ are regulated during early adipogenesis. We identify, in the gene promoters of *Cebpb* and *Cebpd*, the DNA response elements responsible for binding transcription factors that are activated by cAMP or glucocorticoids. We also show that mitogen-activated protein kinase (MAPK)-interacting kinase 2 (MNK2; *Mknk2*), which plays a distinct role in diet-induced obesity, is induced during early adipogenesis and identify the functional DNA response elements responsible for regulating its expression. *Mknk2* expression is maintained in differentiated 3T3-L1 adipocytes and is expressed at high levels across a range of mouse adipose tissue depots. Together, these new insights help to clarify the transcriptional program of early adipogenesis and identify *Mknk2* as one of potentially many genes up-regulated during adipogenesis.

## Introduction

Triacylglycerol (TAG) is a stable and major long-term energy reserve in mammals. TAG stores expand when energy intake exceeds expenditure, leading to individuals becoming overweight or obese; conditions which are associated with a range of chronic disorders including diabetes, cardiovascular disease and some cancers (1,2). Excessive energy promotes weight gain by driving both the expansion of existing adipocytes (hypertrophy) and the development of new ones derived from stem cells (hyperplasia). Given the ever-increasing prevalence of overweight and obesity in the human population, there is an important need to understand the underlying mechanisms that lead to weight gain and diet-induced obesity (DIO), particularly in relation to the development of new adipocytes, a process termed ‘adipogenesis’.

Adipogenesis and the synthesis and breakdown of TAG stores are subject to tight and complex regulation by hormones, neural signals and other pathways (3,4). Established pre-adipocyte cell models have been indispensable for the study of the transcriptional programs that regulate adipogenesis (3,5,6). 3T3-L1 pre-adipocytes are the most widely-used model as they faithfully recapitulate the events that occur during *in vivo* adipogenesis (7–9). Established protocols have been developed to initiate the differentiation of pre-adipocytes; inducers that activate the insulin (10), glucocorticoid and cAMP-signalling pathways (11) initiate a phase of early transcription, which is followed by synchronous entry into the cell cycle and several rounds of mitosis. Upon exit from the cell cycle, cells lose their fibroblastic morphology, accumulate triglyceride and acquire the metabolic features and appearance of adipocytes (7,9–12). TAG accumulation is closely correlated with an increased rate of *de novo* lipogenesis and a coordinate rise in expression of the enzymes of fatty acid and TAG biosynthesis (10,11,13). Rosiglitazone (ROSI), a thiadiazol used as a diabetic treatment to enhance insulin sensitivity (14), is now routinely included in the medium for adipogenic induction as it significantly enhances the conversion of pre-adipocytes to adipocytes (15).

Adipogenesis involves the coordinated and temporal expression of a number of key transcriptional regulators, beginning with the immediate induction of early transcription factors C/EBPβ and C/EBPδ (16). C/EBPβ and C/EBPδ are the direct targets of cAMP- and glucocorticoid signalling, respectively (17, 18). Together, these factors are responsible for inducing the expression of PPARγ which in turn transactivate the expression of C/EBPα. C/EBPα, in concert with PPARγ, co-ordinately activate the expression of adipocyte genes to drive the terminal phase of adipogenesis (19, 20).

The cAMP-elevating agent, 3-isobutyl-1-methylxanthine (IBMX), directly induces expression of *Cebpb*, as mediated by the binding of cAMP response element binding protein (CREB) to a previously identified cAMP response element (CRE) in its promoter (21). *Cebpb* is also reportedly induced by the glucocorticoid dexamethasone (DEX) in hepatocytes (22), muscle (23) and brown adipocytes (24); but this is yet to be shown in differentiating pre-adipocytes. Despite extensive studies of the transcriptional programs regulating early adipogenesis, mapping of the *Cebpb* and *Cebpd* gene promoters (with the exception of (21)) has not been undertaken to identify the CREs and glucocorticoid response elements (GREs) responsible for regulating their expression.

Numerous genes and proteins have been implicated in regulating biological responses to DIO. We recently reported that mice in which the genes for MAPK-interacting kinase 2 (MNK2, *Mknk2*) have been knocked out are protected against DIO (25). Mice in which the genes for the closely-related enzyme, MNK1, are knocked out are also protected against some adverse effects of elevated energy intake (25). Both MNK1 and MNK2 phosphorylate eukaryotic initiation factor 4E (eIF4E), a key component of the cell’s protein synthesis machinery (26). The activity of MNK1 is acutely and markedly stimulated by MAPK signaling, whilst MNK2 has high basal activity that is only slightly elevated by upstream signalling through the MAPKs (27, 28). Whilst regulation of MNK activity is quite well understood, there is no information on the mechanisms that govern their gene expression in differentiating and mature adipocytes, or indeed any other cell type.

In this study, we have probed the fundamental transcriptional events that regulate the expression of C/EBPβ and C/EBPδ. We show that *Cebpb* expression is augmented by DEX and identify the putative GREs likely responsible for mediating this. In addition to the previously reported CRE, we identify an additional distal CRE element that binds CREB in response to elevated cAMP levels. Moreover, we report, for the first time, the presence and location of GREs in the promoter of *Cebpd* which likely regulate its expression. We also investigate how *Mknk2* is transcriptionally regulated during adipogenesis and analyse its expression profile in the adipose depots of mice.

## Materials and Methods

### Cell lines

Murine 3T3-L1 embryonic fibroblasts (ATCC; CL-173) were maintained in high glucose Dulbecco’s modified eagle medium (DMEM) (Invitrogen; 11995-065) supplemented with 10% foetal bovine serum (Invitrogen; 10099-141) and 1% penicillin-streptomycin (Invitrogen; 15140-122) and grown at 37°C in a humidified incubator with 5% CO_2_.

### Differentiation of 3T3-L1 pre-adipocytes

3T3-L1 pre-adipocytes were seeded at a density of 3.5 × 10^4^ cells/cm^2^ and grown to two-days post-confluence (day 0) at which point the media was replaced with growth medium freshly supplemented with 350 nM insulin, 500 *μ*M IBMX, 0.5 *μ*M DEX and 2 *μ*M ROSI (‘differentiation’ media). On day 3, the media was replaced with growth media supplemented with 350 nM insulin (‘maintenance’ media) which was replenished every 3 days.

### RNA isolation

RNA was routinely harvested from cells using TRI Reagent (Sigma; T9424) and cDNA was synthesised using the QuantiNova Reverse Transcription Kit (Qiagen; 205413), according to the manufacturer’s recommendations. Quantitative gene expression analysis was performed using Fast SYBR Green Master Mix (Applied Biosystems; 4385617) on the StepOnePlus Real-Time PCR System (Applied Biosystems, Beverly, MA). RT-qPCR reactions were performed according to the manufacturer’s recommendations using gene-specific primers (**Table 1**). Relative gene expression was calculated using 2^−ΔΔCt^; with the geometric mean of *Nono* and *Hprt* as the internal reference. The relative levels of *Mknk1* and *Mknk2* in mouse tissues were calculated using 2^−ΔCt^ × 100, using the geometric mean of *B2m* and *Hprt* as the internal reference.

### Extraction of protein

Cell monolayers were harvested on ice in RIPA lysis buffer (50 mM Tris-HCl, pH 7.5, 150 mM NaCl, 1% (v/v) NP-40, 1% (v/v) sodium deoxycholate, 0.1% (v/v) sodium dodecyl sulphate (SDS), 1 mM ethylenediaminetetra-acteic acid (EDTA), 50 mM β-glycerophosphate, 0.5 mM NaVO_3_, 0.1% (v/v) 2-mercaptoethanol and 1× protease inhibitors (Roche; 11836170001). Insoluble material was removed by centrifuging at >12,000 × *g* for 10 min at 4°C. Protein content was determined by the Bradford protein assay (Bio-Rad; 5000006) and lysates prepared to equal concentrations.

### Immunoblot analyses

Cell lysates were heated at 95°C for 5 min in Laemmli Sample Buffer and then subjected to sodium-dodecyl sulphate polyacrylamide gel electrophoresis (SDS-PAGE; (52)) followed by electrophoretic transfer to 0.45 *μ*m nitrocellulose membranes (Bio-Rad; 1620115). Membranes were blocked in PBS-0.05% Tween20 (PBST) containing 5% (w/v) skim milk powder for 60 min at room temperature. Membranes were probed with primary antibody in PBST with 5% BSA (w/v) (**Table 2**) overnight at 4°C. Membranes were then washed (3 × 5 min) in PBST and incubated with fluorescently-tagged secondary antibody, diluted in PBST, for 1 h. Membranes were washed again in PBST (3 × 5 min) and fluorescent signals were visualised using the Odyssey Quantitative Imaging System (LI-COR, Lincoln, NE).

### Identification of DNA response element binding sites

Publicly accessible ChIP-sequence (ChIP-seq) data was extracted from ChIP-atlas (chip-atlas.org) using the ‘Peak Browser’ function. Briefly, ChIP-seq data for the antigen of interest was pulled from all cell types using a threshold significance of 100. The data was analysed using the Integrative Genomics Viewer (IGV) software (v2.0) (Broad Institute, Cambridge, MA), aligned to the mouse NCBI37/mm9 genome assembly (July 2007). Enrichment peaks for the antigen of interest were analysed within ± 10 kb of a given gene’s transcription start site. The corresponding regions were analysed on a genome browser (benchling.com) and putative binding sequences were identified using the available consensus sequence data. All reported transcription start site locations are referenced to the mouse NCBI37/mm9 genome assembly (July 2007).

### ChIP

Proteins were cross-linked to DNA by the addition of 37% formaldehyde (Sigma-Aldrich; F8775) to a final concentration of 1% for 10 min at the desired time of analysis. Glycine was then added to a final concentration of 125 mM for 5 min. Cells were washed twice with ice-cold PBS, then collected in 1 mL PBS supplemented with 1× protease inhibitor cocktail and centrifuged at 1000 × *g* for 5 min at 4°C. Cell pellets lysed in 400 μL ChIP lysis buffer (50 mM HEPES-KOH, pH 7.5, 140 mM NaCl, 1mM EDTA pH 8, 1% Triton X-100, 0.1% sodium deoxycholate, 0.1% SDS). Lysates were sonicated on ice for 3 min (in 30 s bursts, with 30 s breaks) after which cell debris was pelleted by centrifugation at 8000 × *g* for 10 min at 4°C. Sonicated chromatin was diluted in 600 μL RIPA buffer (50 mM Tris-HCl, pH 8, 150 mM NaCl, 2 mM EDTA, 1% NP-40, 0.5% sodium deoxycholate, 0.1% SDS), then pre-cleared with 20 *μ*L ChIP-grade Protein G Magnetic Beads (Cell Signaling Technology; 9006) for 2 h at 4°C. 1% of the chromatin was removed to serve as the input control, and then ChIP antibodies (**Table 3**) were added to the chromatin and incubated overnight at 4°C. Immune complexes were collected by adding 20 *μ*L ChIP-grade Protein G Magnetic Beads to each IP for 2 h at 4°C, then washed three times in 1 mL low salt wash buffer (20 mM Tris-HCl, pH 8, 150 mM NaCl, 2 mM EDTA, 1% Triton X-100, 0.1% SDS) for 5 min, followed by a final wash in high salt wash buffer (20 mM Tris-HCl, pH 8, 500 mM NaCl, 2 mM EDTA, 1% Triton X-100, 0.1% SDS). Chromatin was eluted in 150 *μ*L elution buffer (100 mM NaHCO_3_, 1% SDS) for 30 min at 65°C with shaking (1,200 rpm). Cross-links were reversed from eluted chromatin by adding 6 *μ*L of 5 M NaCl and 2 *μ*L Proteinase K (NEB; P8107S) and incubation overnight at 65°C. DNA was purified using QIAquick PCR clean-up kit (Qiagen; 28106) according to the manufacturer’s instructions.

qRT-PCR was performed using 1 *μ*L of eluted chromatin as template with the following cycling conditions: DNA polymerase activation 95°C/180 s, followed by cycles of denaturation (95°C, 15 s) and annealing/extension (60°C, 60 s). Primers were designed manually using Primer3 software to flank the putative DNA binding site(s) for the relevant protein (**Table 4**). Protein enrichment was calculated using the ‘percent input method’ whereby signals obtained from each IP were expressed as a percentage of the total input chromatin.

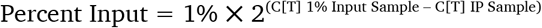

## Results

### Characterisation of rapid changes in gene expression in 3T3-L1 pre-adipocytes

When assessed across the differentiation program we observed that the mRNAs for *Cebpb* and *Cebpd*, two well-established early adipogenic transcription factors, were induced rapidly (3 h after initiating adipogenic induction; Figure 1A). This was subsequently reflected in their respective protein levels (Figure 2A).

**Figure 1.**
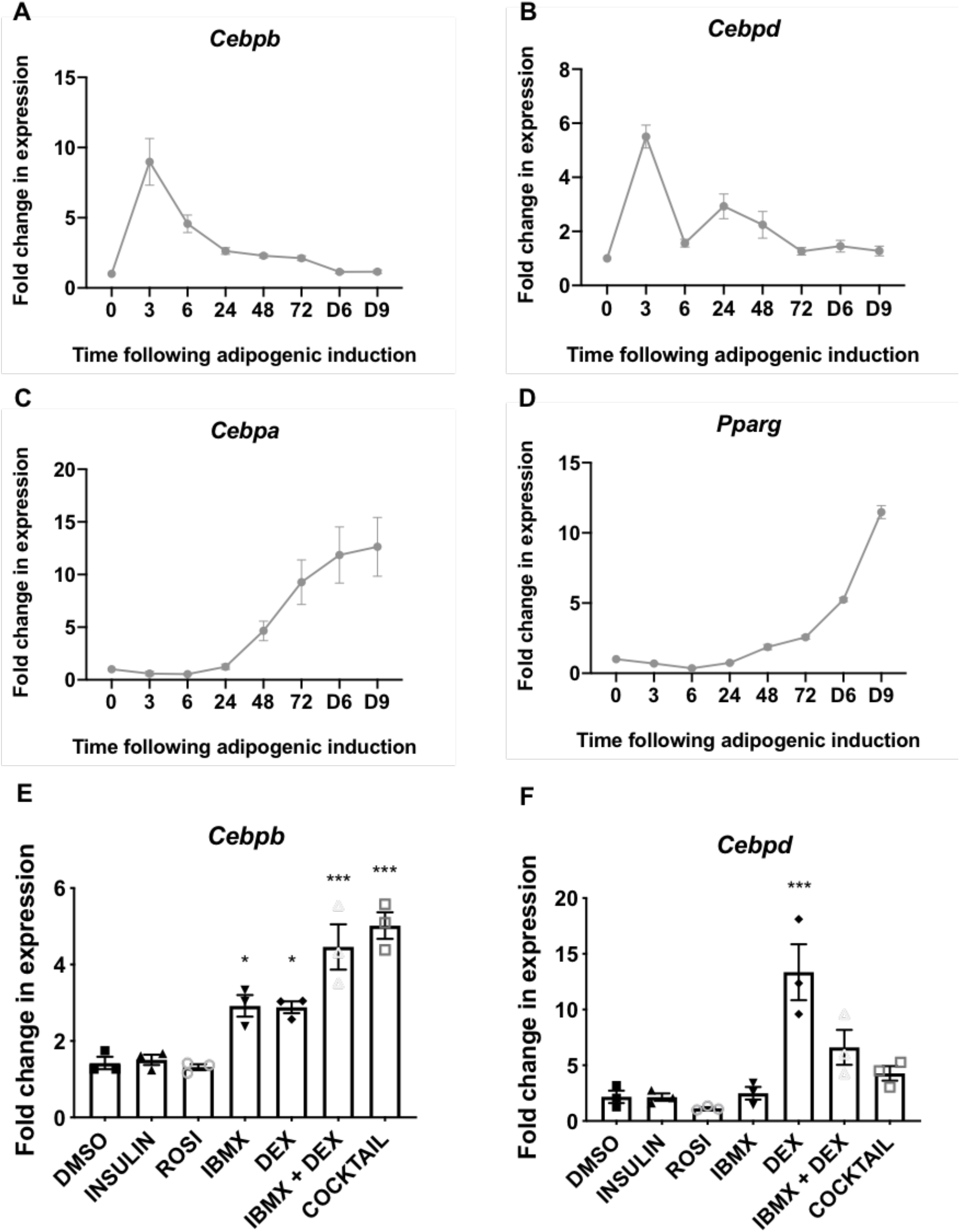
(A-D) 3T3-L1 fibroblasts were treated with the differentiation medium for the indicated times (h, unless indicated by D = days) at which point samples were analysed for expression of *Cebpb, Cebpd, Cebpa* and *Pparg* by RT-qPCR. (E,F) 3T3-L1 fibroblasts were treated with the indicated components of the differentiation medium or the entire cocktail, as indicated, for 3 h at which point samples were analysed for expression of *Cebpb* and *Cebpd* by RT-qPCR. Data presented are mean ± SEM (n=3).

**Figure 2.**
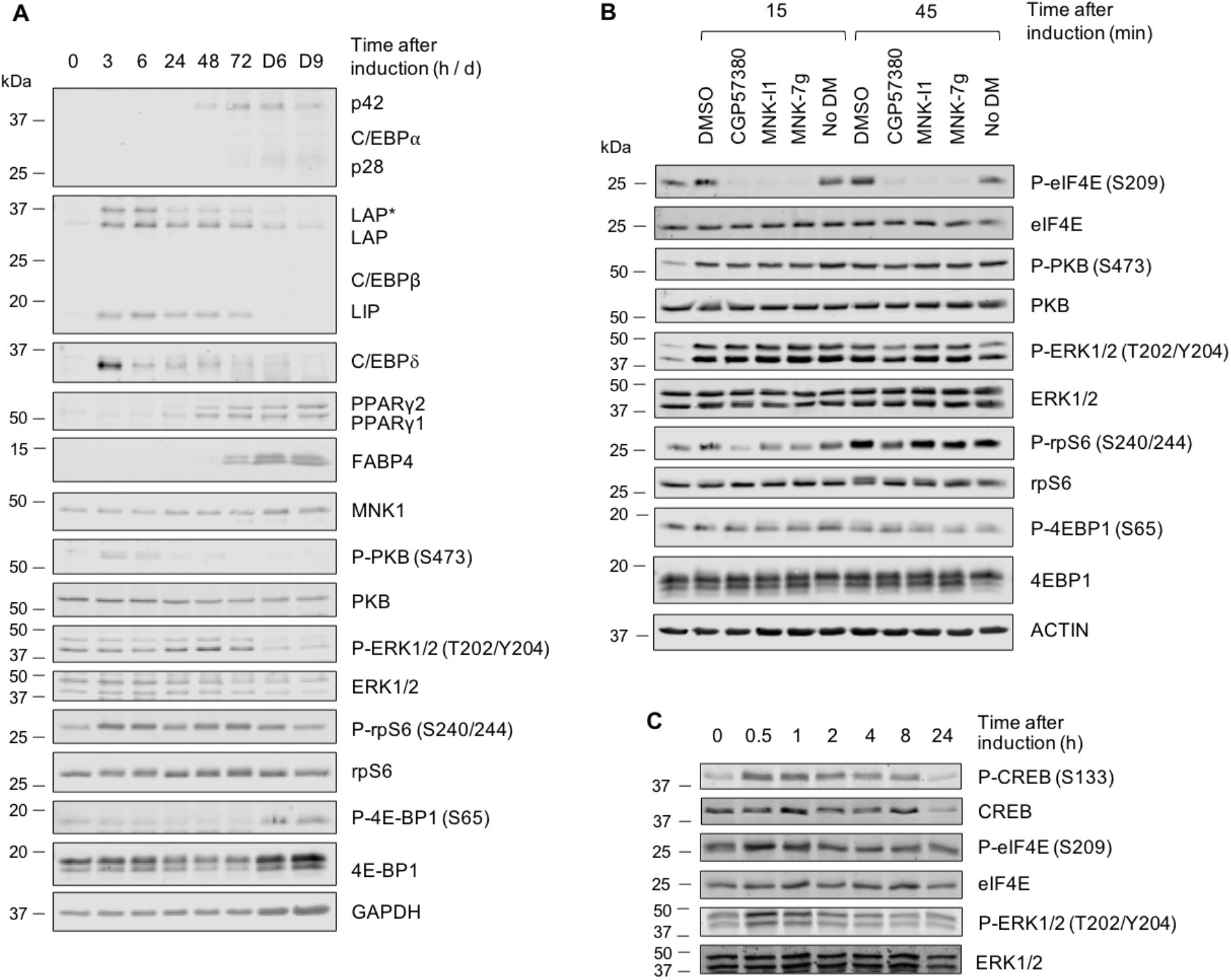
(A-C) 3T3-L1 fibroblasts were treated with differentiation medium for the indicated times (h or min, unless indicated by D = days) at which point cells were harvested and analysed by immunoblot using the indicated antibodies. Data are representative of at least three independent experiments.

Levels of the *Cebpb* and *Cebpd* mRNAs fell thereafter, returning to basal levels by days 3-6 (Figure 2A). Levels of the C/EBPβ and C/EBPδ proteins also gradually declined, but remained above basal levels until at least 72 h after the start of treatment (Figure 2A). *Cebpa* and *Pparg*, encoding the two main drivers of terminal adipogenesis, were induced within 48 h and their mRNA levels continued to rise thereafter (Figure 1C,D). Expression of the C/EBPα and PPARγ proteins increased steadily across the differentiation program, reaching maximal levels in mature, differentiated adipocytes on days 6 and 9 (Figure 2A). PPARγ and C/EBPα drive a program of gene expression that facilitates terminal adipocyte differentiation. One marker of terminal differentiation is fatty acid-binding protein 4 (FABP4), which was induced concomitantly with PPARγ and C/EBPα after 72 h, reaching maximal levels in mature adipocytes on days 6-9 (Figure 2A).

The induction of adipocyte differentiation stimulates several signaling pathways; the level of phosphorylated extracellular signal-regulated kinase (ERK) 1/2 (Thr202/Tyr204) increased after 24 h of adipogenic induction, remained elevated until 72 h but was greatly diminished at days 6 and 9 (Figure 2A). The phosphorylation of ribosomal protein S6 (rpS6) (Ser240/244), a readout of mTORC1 signaling, was elevated at 3, 6, 48 and 72 h (Figure 2A). Lower P-rpS6 levels however, were recorded at 24 h and on days 6 and 9. Total levels of eIF4E-binding protein 1 (4E-BP1), the availability and phosphorylation of which regulates the initiation of mRNA translation, were elevated in terminally differentiated adipocytes on days 6 and 9. However, the phosphorylation of 4E-BP1 (P-4E-BP1), which is catalysed by mTORC1, was not increased relative to total levels at those times (Figure 2A). The phosphorylation of protein kinase B (PKB, also termed Akt) (Ser473), which is a marker of phosphoinositide 3-kinase (PI3K)/insulin signalling was acutely elevated at 3 and 6 h, but this fell by 24 h. Analysis at earlier time points (15 and 45 min) revealed that the differentiation induction rapidly activated the ERK and PKB pathways, while mTORC1 signaling lagged somewhat, only being evident by 45 min (Figure 2B). Thus, the adipogenic cocktail induces rapid but transient activation of PKB, swift and sustained activation of the mTORC1 pathway and slower stimulation of the ERK MAPK pathway which is sustained up to 72 h.

The differentiation medium (which contains the cAMP-phosphodiesterase inhibitor IBMX) rapidly stimulated the phosphorylation of CREB, a transcription factor which is the direct substrate for cAMP-dependent protein kinase, PKA (Figure 2C). Its increased phosphorylation was already evident at 30 min and remained elevated until 1 h, after which it gradually declined. We also observed modest and transient rises in the phosphorylation of the MNK substrate eIF4E and of ERK1/2, which lie upstream of the MNKs (Figure 2C). Since MNK1, in particular, is activated by ERK (26,29) its activation likely explains the transient increase in P-eIF4E while the basal level of P-eIF4E is likely due to MNK2, which has constitutive activity (28).

### Induction of early gene expression by individual components of the adipogenic cocktail

As shown above, adipogenic stimulation resulted in an immediate induction in *Cebpb* and *Cebpd*. To understand how early adipogenic signaling brings about this induction, individual components of the adipogenic induction medium were tested to assess their roles and dissect the signalling pathways involved. By activating PKA, IBMX induces the phosphorylation of CREB, which then enters the nucleus and binds to cAMP response elements (CREs) to activate expression of target genes. DEX is a glucocorticoid receptor (GR) agonist, which binds to and activates the GR. Ligand-bound GR translocates to the nucleus where it binds to glucocorticoid response elements (GREs) to promote the expression of its target genes. Insulin is an anabolic hormone that binds to its extracellular receptor which in turn activates PI3K/PKB signalling and leads to diverse cellular responses. ROSI is a synthetic ligand that promotes PPARγ activity during terminal adipogenesis and (because PPARγ is only present later in adipogenesis) is not expected to directly modulate early adipogenic signalling. These components were used either individually, in combination or all together (‘cocktail’) to assess their abilities to induce *Cebpb, Cebpd, Mknk1* and *Mknk2* during early adipogenesis.

IBMX and DEX each induced similar levels of *Cebpb* mRNA expression and its levels were further increased when IBMX and DEX were used in combination (Figure 1E). *Cebpb* expression was not further elevated by the addition of insulin and ROSI, suggesting IBMX and DEX are responsible for inducing *Cebpb* expression through separate pathways (Figure 1E).

DEX alone induced *Cebpd* (Figure 1F), while IBMX alone did not; rather, when combined with DEX, IBMX actually reduced expression to half the level observed with DEX alone (Figure 1F). The addition of insulin and ROSI further blunted *Cebpd* expression, suggesting DEX alone is responsible for *Cebpd* expression, and that the effect of DEX on *Cebpd* is negatively regulated by IBMX, insulin and/or ROSI.

### Transcription factor binding sites in the *Cebpb* and *Cebpd* genes

Apart from a study mapping the CRE-like elements responsible for cAMP-induced *Cebpb* expression (21) and a genome-wide study identifying putative CRE elements based on computational prediction (30) there have been no detailed studies mapping transcription factor binding sites on the promoters of *Cebpb* or *Cebpd, Mknk1 or Mknk2*. Publicly-accessible ChIP-seq data were therefore used to locate possible CRE’s and GRE’s, respectively, in these genes’ promoters. Analysis identified three GRE-like elements in *Cebpb*, two of which were proximal (−1161 and −2370) to the transcription start site (TSS), the other being located in the distal promoter (+20) (Figure 3A). The previously-mapped CRE-like element in *Cebpb* was identified (−60/106) (21), along with two additional CRE-like elements (−2343/2398) (Figure 3-3A). The distal CRE-like elements identified in this study shared the half-CRE core sequence (TGACG), with the exception of their final nucleotide (TGAC**A**/ TGAC**C**) (Figure 3A).

**Figure 3.**
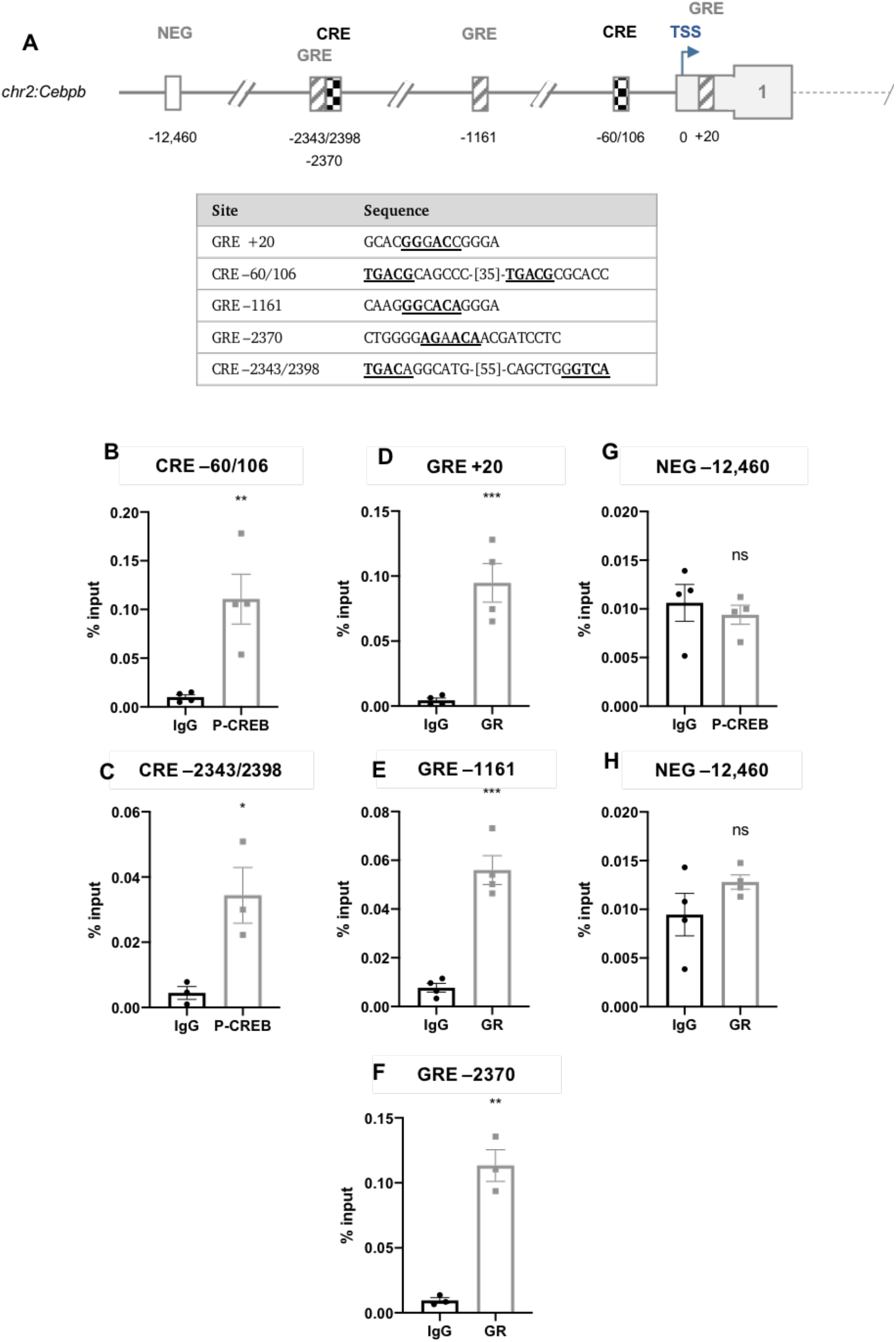
(A) Schematic diagram depicting the genomic position of putative response elements on the *Cebpb* promoter relative to the TSS (chr2:167,514,466). The sequences of the putative response elements are displayed in the table; nucleotides corresponding to the core consensus sequence are shown bold and underlined. ChIP assays were performed in 3T3-L1 fibroblasts to assess enrichment of (B,C) P-CREB at putative CREs and (D-F) GR at putative GREs after 3 h of adipogenic induction. (G,H) Enrichment was also assessed at the indicated NEG region. Data are mean ± SEM; n=4; unpaired two-tailed t-test (*P < 0.05 **P < 0.01 ***P < 0.001). Abbreviations: GRE: glucocorticoid response element; CRE: cAMP response element; NEG: negative non-specific region; TSS: transcription start site.

In *Cebpd*, only a single CRE- (−41) and GRE-like element (−98) were identified in the proximal promoter (Figure 4A). The CRE (−41) shared the half-CRE core sequence (Figure 4A). There are no published reports of CRE or GRE features in the promoter region of the *Cebpd* gene.

**Figure 4.**
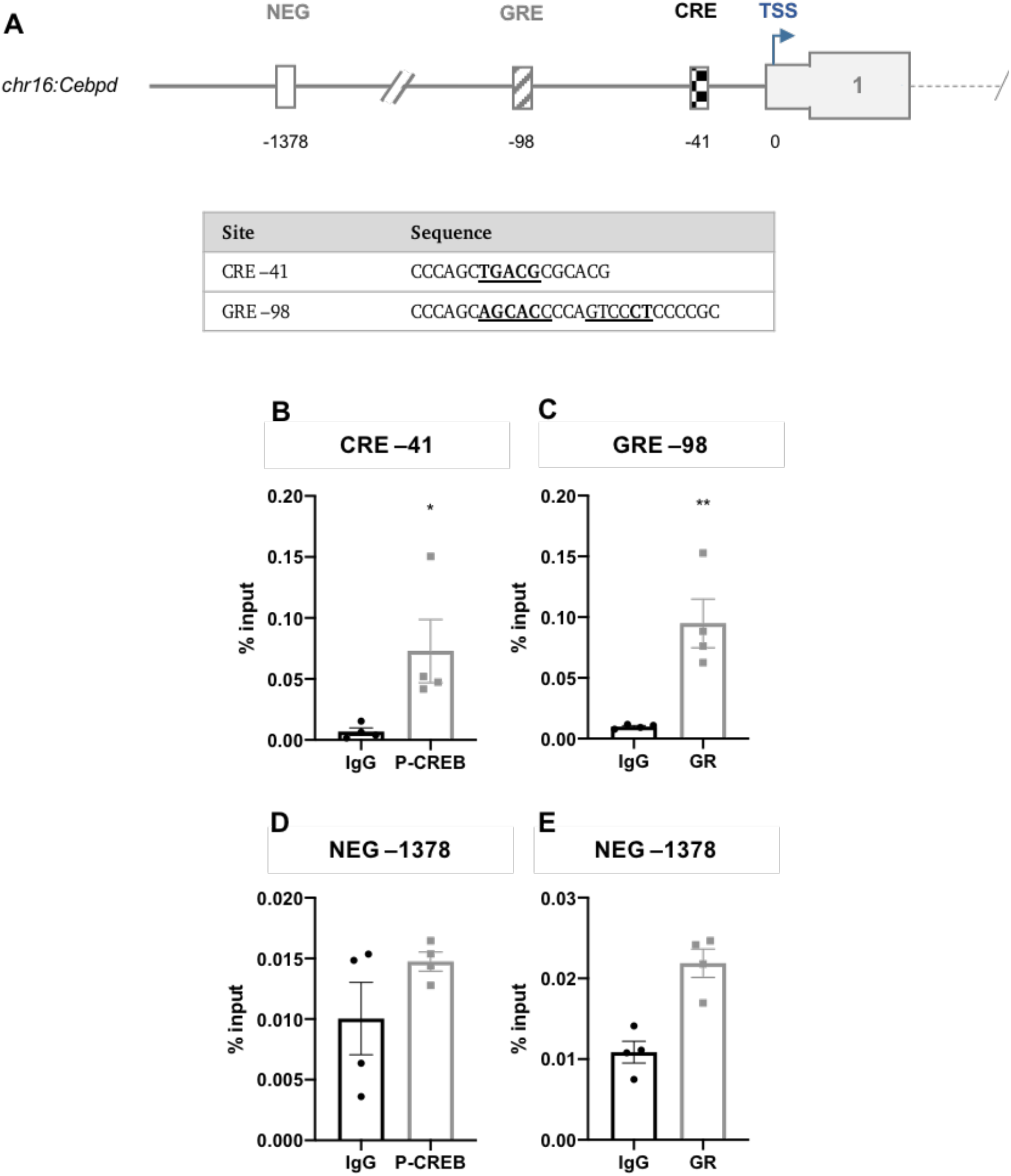
(A) Schematic diagram depicting the genomic position of putative response elements on the *Cebpd* promoter relative to the TSS (chr16:15,887,379). The sequences of the putative response elements are displayed in the table; nucleotides corresponding to the core consensus sequence are shown bold and underlined. ChIP assays were performed in 3T3-L1 fibroblasts to assess enrichment of (B) P-CREB at putative CREs and (C) GR at putative GREs after 3 h of adipogenic induction. (D,E) Enrichment was also assessed at the indicated NEG region. Data are mean ± SEM; n=4; unpaired two-tailed t-test (*P < 0.05 **P < 0.01 ***P < 0.001). Abbreviations: GRE: glucocorticoid response element; CRE: cAMP response element; NEG: negative non-specific region; TSS: transcription start site.

ChIP assays were performed to enrich for CREB and GR at the CRE- and GRE-like elements following 3 h of adipogenic stimulation. Antibodies for P-CREB (Ser133) and GR were used for ChIP. In *Cebpb*, P-CREB was most significantly enriched at the −60/106 CRE, consistent with the findings of Zhang *et al.* (21) which identified a CRE in this region, and also at the −2343/2398 CRE (Figure 3B,C). GR was most significantly enriched at the +20 and −2370 GRE-like elements but less significantly at the −1161 element (Figure 3D-F). In *Cebpd*, P-CREB and GR were significantly enriched at the −41 CRE- and −98 GRE-like elements, respectively (Figure 4B,C). There was no significant enrichment in the negative non-specific (NEG) controls indicating that enrichment observed at the proposed GREs was specific to the proposed response elements (Figures 3G,H and 4D,E).

### Control of expression of the MNKs

We have previously shown that mice in which the genes for MNK1 (*Mknk1*) or MNK2 (*Mknk2*) are knocked out are protected against DIO and provided data suggesting that MNKs may regulate adipogenesis (25). However, there is no information concerning the regulation of the expression of MNK1 or MNK2 or the transcription factors that may be involved.

Analysis of the expression of the *Mknk1* and *Mknk2* genes in different mouse tissues revealed that theyare expressed at relatively low levels in liver and subcutaneous fat (Figure 5A,B). In omental, gonadal and scapular adipose tissue and especially in muscle, *Mknk2* is highly expressed and appears to be the predominant MNK isoform (Figure 5A,B). Interestingly, *Mknk2* appears to be expressed at somewhat lower levels on a high-fat diet, most evidently in scapular adipose tissue (Figure 5B).

**Figure 5.**
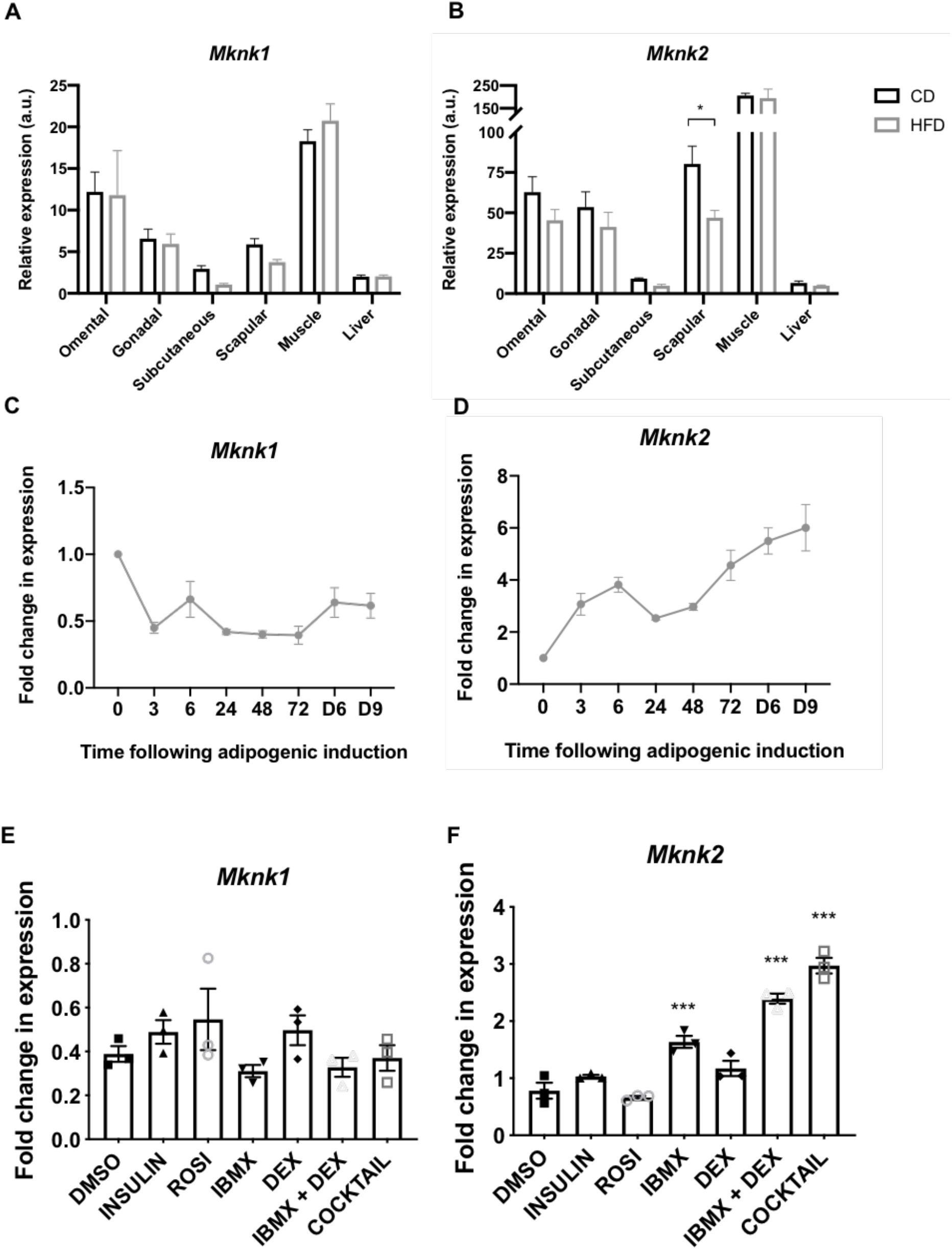
(A,B) RT-qPCR analysis of *Mknk1* and *Mknk2* expression in the indicated tissues of C57BL6/J mice fed chow or a high fat diet for 16 weeks. Data presented are mean ± SEM (n=5-6 per group). (C,D) 3T3-L1 fibroblasts were treated with the differentiation medium for the indicated times (h, unless denoted by D = days) at which point samples were analysed for expression of *Mknk1* and *Mknk2* by RT-qPCR. (E,F) 3T3-L1 fibroblasts were treated with the indicated components of the differentiation medium or the entire cocktail, as indicated, for 3 h at which point samples were analysed for expression of *Mknk1* and *Mknk2* by RT-qPCR. Data presented are mean ± SEM (n=3).

Given the role of the MNKs in 3T3-L1 differentiation, it was of interest to examine whether their expression changed during adipogenesis. The expression of *Mknk1* declined within 3 h but then remained steady (Figure 5C). In contrast, *Mknk2* mRNA levels increased quickly after addition of the cocktail and continued to increase up to day 9 where it was maintained in mature adipocytes (Figure 5D).

Analysis of the effects of individual components of the cocktail showed that no individual agent or any combination tested significantly altered the levels of *Mknk1* mRNA (Figure 5E). IBMX alone increased *Mknk2* mRNA levels and this effect was enhanced by co-treatment with DEX and further by the full cocktail (Figure 5F). This suggests that cAMP and glucocorticoid signaling regulate the expression of *Mknk2*.

In *Mknk1*, a single CRE- (−60) and GRE-like element (−142) were identified (Figure 6A). A single CRE-like element was identified in *Mknk2* (−372) along with three GRE-like elements (−86, −1883, −3270) (Figure 6B). The CRE-like elements in both *Mknk1* (TGAC**T**)and *Mknk2* (TGAC**C**) share the half-CRE core sequence with the exception of the final nucleotide, as indicated in bold.

**Figure 6.**
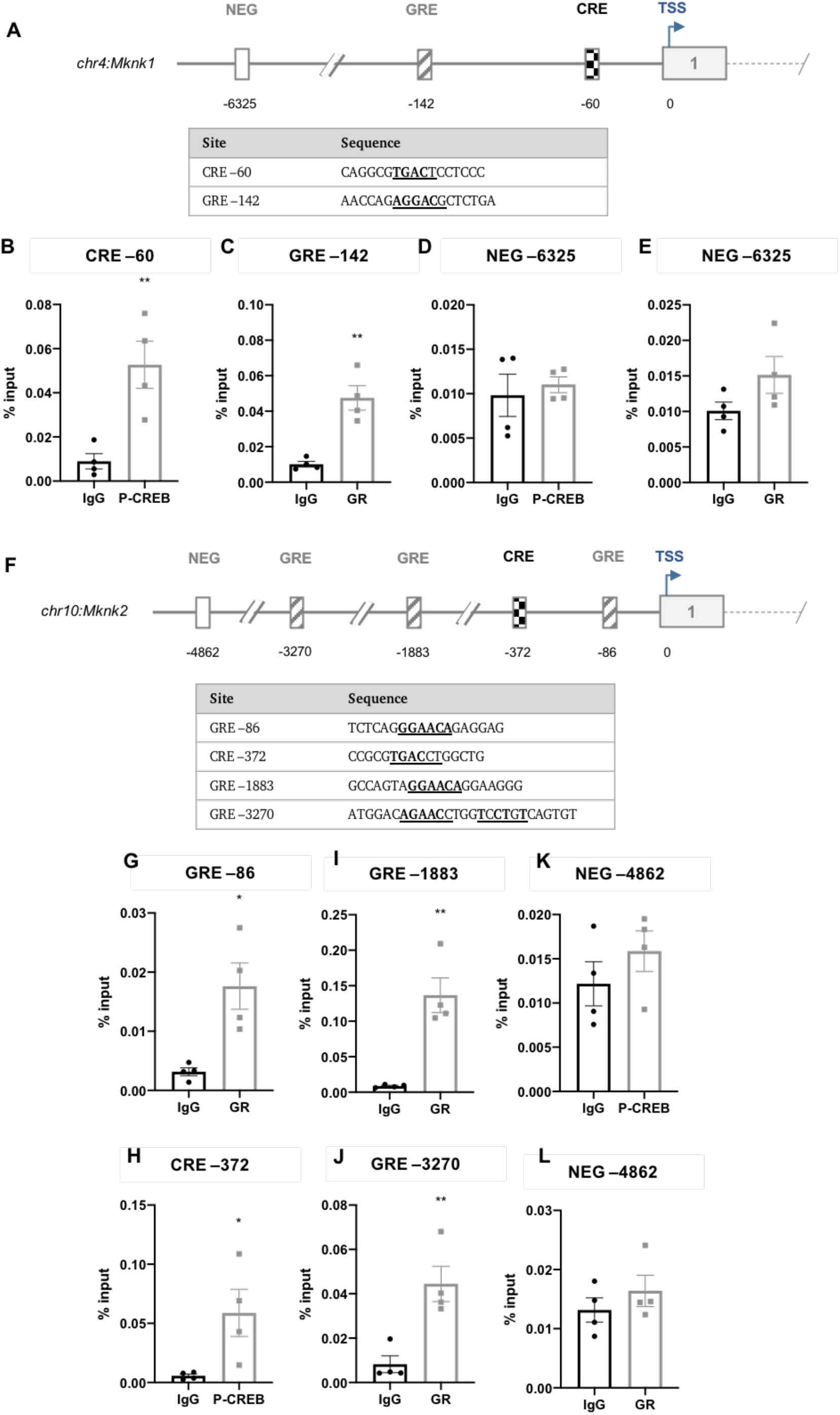
Schematic diagram depicting the genomic position of putative response elements on the (A) *Mknk1* and (F) *Mknk2* promoters relative to their TSS (chr4:115,511,826 and chr10:80,139,038, respectively). The sequences of the putative response elements are displayed in the tables below; nucleotides corresponding to the core consensus sequence are shown bold and underlined. ChIP assays were performed in 3T3-L1 fibroblasts to assess enrichment of (B,H) P-CREB at putative CREs and (C,G,I,J) GR at putative GREs after 3 h of adipogenic induction. (D,E,K,L) Enrichment was also assessed at the indicated NEG regions of the two genes. Data are mean ± SEM; n=4; unpaired two-tailed t-test (*P < 0.05 **P < 0.01 ***P < 0.001). Abbreviations: GRE: glucocorticoid response element; CRE: cAMP response element; NEG: negative non-specific region; TSS: transcription start site.

ChIP analysis using antibodies for P-CREB or the GR revealed that P-CREB and the GR were significantly enriched in *Mknk1* at the −60 CRE and −142 GRE-like elements, respectively (Figure 6B,C). P-CREB was significantly enriched at the *Mknk2* CRE (−356) (Figure 6H), whilst GR was most significantly enriched at the −1883 GRE but less so at the −86 and −3385 GREs (Figure 6G,I,J). Enrichment was specific to the putative response elements as there was no significant enrichment in the negative non-specific (NEG) regions (Figures 6D,E,K,L).

Given that *Mknk2* is up-regulated substantially and in a sustained manner during adipogenesis, it seemed possible that adipogenic transcription factors were involved in up-regulating its expression. Again, publicly-accessible ChIP-seq data were used to analyse the enrichment peaks of C/EBPα, C/EBPβ and PPARγ to locate putative C/EBP response elements (C/EBPREs) and PPAR response elements (PPAREs) in the promoter of *Mknk2.* The analysis identified two putative C/EBPREs (−264/290 and −1204/1258) and three putative PPAREs (−1975, −3348 and −3993) (Figure 7A). When ChIP assays were performed to assess enrichment of PPARγ after 72 h there was no significant enrichment at any of the PPARE elements (Figure 7B-D). ChIP assays were performed to enrich for C/EBPβ and C/EBPα at the C/EBPRE elements following 24 and 72 h of adipogenic induction, respectively, accounting for the temporal separation in their expression. C/EBPβ was enriched at the −1204/1258 region but less so at the −264/290 C/EBPRE, although neither site was enriched significantly (Figure 7E,F). The ChIP-seq analysis predicted C/EBPα occupancy at the −1204/1258 but not the −264/290 C/EBPRE; however, no significant enrichment of C/EBPα was observed at the −1204/1258 element (Figure 7G). Enrichment was specific to the putative response elements as there was no significant enrichment in the negative non-specific (NEG) regions (Figures 7H-J).

**Figure 7.**
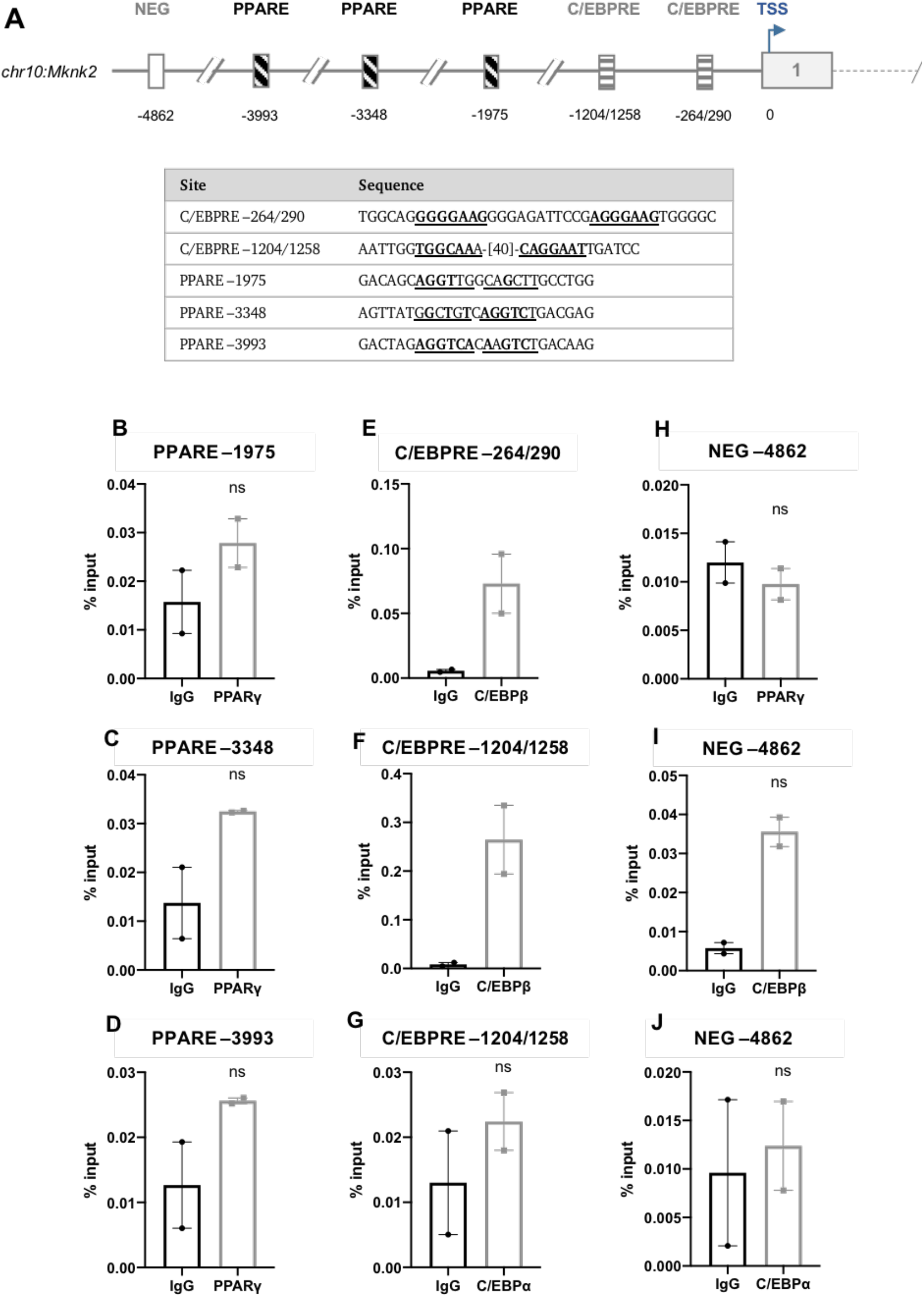
(A) Schematic diagram depicting the genomic position of putative response elements on th *Mknk2* promoter relative to the TSS (chr10:80,139,038). The sequences of the putative response elements are displayed; nucleotides corresponding to the core consensus sequence are shown bold and underlined. ChIP assays were performed in 3T3-L1 fibroblasts to assess enrichment of (B-D) PPARγ at putative PPAREs and (G) C/EBPα at putative C/EBPREs after 72 h of induction. (E,F) Enrichment of C/EBPβ at putative C/EBPREs was assessed after 24 h. (H-J) Enrichment was also assessed at the indicated NEG region. Data are mean ± SEM; n=2; unpaired two-tailed t-test (*P < 0.05 **P < 0.01 ***P < 0.001). Abbreviations: PPARE: PPAR response element; C/EBPRE: C/EBP response element; NEG: negative non-specific region; TSS: transcription start site.

## Discussion

In this study, we investigated the early transcriptional program of adipogenesis and revealed how the expression of two important adipogenic transcription factors, C/EBPβ and C/EBPδ, are regulated in 3T3-L1 pre-adipocytes. We identified (or refined) for the first time, the DNA response elements in the promoters of *Cebpb* and *Cebpd* that are bound by transcription factors activated by cAMP and glucocorticoids. We have analysed the expression of MNK1/2 during 3T3-L1 adipogenesis and identified the response elements likely responsible for regulating their expression. We also showed that *Mknk2* is the predominant isoform in mouse adipose tissue and that it may be changed when mice consume a high-fat diet.

C/EBPβ and C/EBPδ have important and partially redundant roles in adipogenesis, in part by activating expression of PPARγ (17,31–34). It is well-established that IBMX directly induces expression of *Cebpb*, as mediated by the binding of CREB to a previously identified CRE (−60/106) (21). Upon analysis of the *Cebpb* promoter, however, we located an additional CRE-like element at −2343/2398 which was occupied by CREB, albeit to a lesser extent (Figure 3B,C). This suggests that CREB may regulate *Cebpb* expression from more than one CRE. However, Zhang *et al.* (21) did not identify such a distal CRE-like element and their gene-reporter constructs indicated that sequences greater 100 nucleotides from the TSS were not required for reporter expression. Therefore, the functional significance of the −2343/2398 site remains unknown until determined experimentally using a similar gene-reporter assay.

Notably, and for the first time, DEX was found to directly induce the expression of *Cebpb* (Figure 1E) through the occupancy of GR at three half-GRE core sequence elements in its proximal and distal promoter (Figure 3). Previous reports of DEX-induced *Cebpb* expression used other cell systems (22–24) and this is the first report of DEX-induced *Cebpb* expression in pre-adipocytes. Furthermore, when DEX was combined with IBMX, it was found to further augment *Cebpb* expression (Figure 1E). Augmented *Cebpb* induction has been previously reported in hepatocytes using DEX in combination with either glucagon (a beta-adrenergic receptor agonist that increases cAMP) (35) or a cAMP analogue (36). The combination of DEX and glucagon was shown to facilitate a greater maximal increase and more gradual decline in *Cebpb* levels than was observed with either DEX or glucagon alone (35). This suggests that IBMX and DEX are each necessary for maximal and sustained C/EBPβ expression and may help to explain why pre-adipocytes incubated in the absence of DEX show diminished expression of C/EBPβ, C/EBPδ and C/EBPα and very little evidence of differentiation (17). These findings challenge the existing adipogenic transcriptional paradigm which assumes that *Cebpb* is solely induced by IBMX (17,18). It may also have implications for other IBMX-driven genes and suggests a complex relationship between CREB and GR-driven gene expression.

It is well-established that DEX directly induces the expression of *Cebpd* (17,18,23) but the location of the responsible GRE(s) has not been previously reported. In this study, two imperfect palindrome 6-bp half-sites separated by three nucleotides were identified in the *Cebpd* proximal promoter (−98), corresponding to a full-length GRE consensus sequence (Figure 4A). This site was found to be occupied by the GR following adipogenic induction, and is therefore likely to be the GRE element responsible for regulating *Cebpd* expression (Figure 4C). Analysis also identified a half-CRE element in the *Cebpd* promoter (Figure 4B); however, unlike *Cebpb*, IBMX and DEX did not augment *Cebpd* expression. Rather, IBMX actually impaired *Cebpd* expression suggesting the CRE-like element may somehow repress *Cebpd* expression (Figure 1F). Given that IBMX is present in the initial induction media, such down-regulation of *Cebpd* appears counter-intuitive; however, this may be a mechanism by which cells appropriately modulate expression of *Cebpd* given that C/EBPδ is already very highly expressed after 3 h in the presence of IBMX (Figure 2A). CREB is typically thought of as a factor that activates gene expression but it can also act to negatively regulate gene expression (37–40). The regulation of CREB-mediated transcription appears to be gene- and context-specific dependent on the recruitment of co-activators and co-repressors which vary between promoter environments (41, 42). CREB may therefore negatively regulate *Cebpd* levels to modulate and balance its expression through recruitment of co-repressors at or near its promoter; however, further work is required to investigate this.

Here it was shown that *Mknk2*, but not *Mknk1*, is progressively increased during adipogenesis (Figure 5D). In the first analysis of the *Mknk2* promoter, we identified multiple GRE-like elements bound by GR and a single CRE-like element bound by CREB, indicating that *Mknk2* expression is likely regulated through cAMP and GR signalling (Figure 6F-J). During adipogenesis, such pathways are activated immediately upon hormone induction (17,18), suggesting MNK2 up-regulation may be important for adipocyte differentiation. These findings suggest *Mknk2* may be just one of a number of other genes up-regulated by CREB and GR to promote adipogenesis (24, 43). Indeed, given its expression in mature adipocytes (Figure 5B), MNK2 may well perform other functions that regulate adipocyte biology. On another note, the finding that *Mknk2* is regulated by cAMP and glucocorticoids may provide a useful insight into certain types of cancers in which MNK2 is highly expressed and associated with poor prognosis (44–46).

In contrast, *Mknk1* levels remained quite stable (Figure 5C), despite the occupancy of its putative CRE and GRE-like elements by CREB and GR, respectively (Figure 6A-C). However, it is possible that whilst CREB occupancy on the CRE of *Mknk2* is inducible (47), the CRE-like element in *Mknk1* may be constitutively occupied, as has been observed in the gene promoters for *Pepck*, *Pcna*, *Ccna* and *c-Fos* (48–51). Based upon their differential regulation, MNK1/2 may therefore perform distinct functions in adipocytes. The expression levels of *Mknk1* and *Mknk2* in differentiated 3T3-L1 adipocytes was reflected in the expression profile of mouse adipose tissue; *Mknk2* was highly expressed and predominated over *Mknk1* in omental, gonadal and scapular adipose tissue (Figure 5A,B). The expression of *Mknk2* in gonadal fat was consistent with earlier findings (25), and suggests more generally that MNK2 may be important for regulating aspects of adipocyte biology. The expression of *Mknk1* and *Mknk2* was relatively low in subcutaneous adipose tissue (Figure 5A,B), suggesting there may be a lesser role for MNKs in this depot. *Mknk2* expression was lower in adipose tissue from mice given a high-fat diet, most evidently in scapular adipose tissue (Figure 5B). This suggests that *Mknk2* expression responds to changes in diet, perhaps allowing it to modulate changes in adipocyte metabolism. This may help to explain why MNK2-KO mice are protected from DIO (25), although the detailed mechanisms remain to be established.

In conclusion, these studies provide new insights on the elements that control the expression of *Cebpb* and *Cebpd* during early adipogenesis. We also provide the first report, in any species or system, about the transcriptional regulation of *Mknk2* and allude to a potential role for MNK2 in adipocyte biology.

## Acknowledgments

The 3T3-L1 fibroblasts used in this study were generously provided by Yeeshim Khew-Goodhall from the Centre for Cancer Biology (SA Pathology, Adelaide, South Australia).

## Funding details

This work was supported by funding from the South Australian Health and Medical Research Institute. JEM acknowledges the support received through the provision of an Australian Government Research Training Program Scholarship. PJP is supported by a L2 Future Leader Fellowship from the National Heart Foundation of Australia [FLF102056] and L2 Career Development Fellowship from the National Health and Medical Research Council [CDF1161506].

## Declaration of interest statement

The authors declare that they have no conflicts of interest with the contents of this article.

## Notes

### Competing Interest Statement

The authors have declared no competing interest.

## References

1. O’Neill, S., and O’Driscoll, L. (2014) Metabolic syndrome: a closer look at the growing epidemic and its associated pathologies. Obesity Reviews. 16, 1–12

2. Upadhyay, J., Farr, O., Perakakis, N., Ghaly, W., and Mantzoros, C. (2018) Obesity as a Disease. Medical Clinics of North America. 102, 13–33.

3. Tang, Q. Q., and Lane, M. D. (2012) Adipogenesis: From Stem Cell to Adipocyte. Annual Review of Biochemistry. 81, 715–736.

4. Pellegrinelli, V., Carobbio, S., and Vidal-Puig, A. (2016) Adipose tissue plasticity: how fat depots respond differently to pathophysiological cues. Diabetologia. 59, 1075–1088.

5. Farmer, S. R. (2006) Transcriptional control of adipocyte formation. Cell Metabolism. 4, 263–273.

6. Lowe, C. E., O’Rahilly, S., and Rochford, J. J. (2011) Adipogenesis at a glance. Journal of Cell Science. 124, 3726–3726.

7. Green, H., and Kehinde, O. (1975) An established preadipose cell line and its differentiation in culture II. Factors affecting the adipose conversion. Cell. 5, 19–27.

8. Green, H., and Kehinde, O. (1976) Spontaneous heritable changes leading to increased adipose conversion in 3T3 cells. Cell. 7, 105–113.

9. Green, H., and Meuth, M. (1974) An established pre-adipose cell line and its differentiation in culture. Cell. 3, 127–133.

10. MacDougald, O. A., and Lane, M. D. (1995) Transcriptional Regulation of Gene Expression During Adipocyte Differentiation. Annual Review of Biochemistry. 64, 345–373.

11. Student, A. K., Hsu, R. Y., and Lane, M. D. (1980) Induction of fatty acid synthetase synthesis in differentiating 3T3-L1 preadipocytes. The Journal of biological chemistry. 255, 4745–50.

12. Green, H., and Kehinde, O. (1974) Sublines of mouse 3T3 cells that accumulate lipid. Cell. 1, 113–116.

13. Coleman, R. A., Reed, B. C., Mackall, J. C., Student, A. K., Lane, M. D., and Bell, R. M. (1978) Selective changes in microsomal enzymes of triacylglycerol phosphatidylcholine, and phosphatidylethanolamine biosynthesis during differentiation of 3T3-L1 preadipocytes. The Journal of biological chemistry. 253, 7256–61.

14. Day, C. (1999) Thiazolidinediones: a new class of antidiabetic drugs. Diabetic Medicine. 16, 179–192.

15. Zebisch, K., Voigt, V., Wabitsch, M., and Brandsch, M. (2012) Protocol for effective differentiation of 3T3-L1 cells to adipocytes. Analytical Biochemistry. 425, 88–90.

16. Tang, Q.-Q., and Lane, M. D. (1999) Activation and centromeric localization of CCAAT/enhancer-binding proteins during the mitotic clonal expansion of adipocyte differentiation. Genes & Development. 13, 2231–2241

17. Yeh, W. C., Cao, Z., Classon, M., and McKnight, S. L. (1995) Cascade regulation of terminal adipocyte differentiation by three members of the C/EBP family of leucine zipper proteins. Genes & Development. 9, 168–181

18. Cao, Z., Umek, R. M., and McKnight, S. L. (1991) Regulated expression of three C/EBP isoforms during adipose conversion of 3T3-L1 cells. Genes & Development. 5, 1538–1552.

19. Hamm, J. K., Park, B. H., and Farmer, S. R. (2001) A Role for C/EBPβ in Regulating Peroxisome Proliferator-activated Receptor γ Activity during Adipogenesis in 3T3-L1 Preadipocytes. Journal of Biological Chemistry. 276, 18464–18471.

20. Wu, Z., Rosen, E. D., Brun, R., Hauser, S., Adelmant, G., Troy, A. E., McKeon, C., Darlington, G. J., and Spiegelman, B. M. (1999) Cross-Regulation of C/EBPα and PPARγ Controls the Transcriptional Pathway of Adipogenesis and Insulin Sensitivity. Molecular Cell. 3, 151–158.

21. Zhang, J.-W., Klemm, D. J., Vinson, C., and Lane, M. D. (2004) Role of CREB in Transcriptional Regulation of CCAAT/Enhancer-binding Protein β Gene during Adipogenesis. Journal of Biological Chemistry. 279, 4471–4478.

22. Campos, S. P., and Baumann, H. (1992) Insulin is a prominent modulator of the cytokine-stimulated expression of acute-phase plasma protein genes. Molecular and Cellular Biology. 12, 1789–1797.

23. Yang, H., Mammen, J., Wei, W., Menconi, M., Evenson, A., Fareed, M., Petkova, V., and Hasselgren, P. (2005) Expression and activity of C/EBPβ and δ are upregulated by dexamethasone in skeletal muscle. Journal of Cellular Physiology. 204, 219–226.

24. Park, Y.-K., and Ge, K. (2016) Glucocorticoid Receptor Accelerates, but Is Dispensable for, Adipogenesis. Molecular and Cellular Biology. 10.1128/mcb.00260-16

25. Moore, C. E. J., Pickford, J., Cagampang, F. R., Stead, R. L., Tian, S., Zhao, X., Tang, X., Byrne, C. D., and Proud, C. G. (2016) MNK1 and MNK2 mediate adverse effects of high-fat feeding in distinct ways. Scientific Reports. 10.1038/srep23476

26. Waskiewicz, A. J., Flynn, A., Proud, C. G., and Cooper, J. A. (1997) Mitogen-activated protein kinases activate the serine/threonine kinases Mnk1 and Mnk2. The EMBO Journal. 16, 1909–1920.

27. Buxade, M., Parra-Palau, J. L., and Proud, C. G. (2008) The Mnks: MAP kinase-interacting kinases (MAP kinase signal-integrating kinases). Frontiers in Bioscience. Volume, 5359

28. Scheper, G. C., Morrice, N. A., Kleijn, M., and Proud, C. G. (2001) The Mitogen-Activated Protein Kinase Signal-Integrating Kinase Mnk2 Is a Eukaryotic Initiation Factor 4E Kinase with High Levels of Basal Activity in Mammalian Cells. Molecular and Cellular Biology. 21, 743–754.

29. Wang, X., Flynn, A., Waskiewicz, A. J., Webb, B. L. J., Vries, R. G., Baines, I. A., Cooper, J. A., and Proud, C. G. (1998) The Phosphorylation of Eukaryotic Initiation Factor eIF4E in Response to Phorbol Esters, Cell Stresses, and Cytokines Is Mediated by Distinct MAP Kinase Pathways. Journal of Biological Chemistry. 273, 9373–9377.

30. Zhang, X., Odom, D. T., Koo, S.-H., Conkright, M. D., Canettieri, G., Best, J., Chen, H., Jenner, R., Herbolsheimer, E., Jacobsen, E., Kadam, S., Ecker, J. R., Emerson, B., Hogenesch, J. B., Unterman, T., Young, R. A., and Montminy, M. (2005) Genome-wide analysis of cAMP-response element binding protein occupancy, phosphorylation, and target gene activation in human tissues. Proceedings of the National Academy of Sciences. 102, 4459–4464.

31. Tanaka, T., Yoshida, N., Kishimoto, T., and Akira, S. (1997) Defective adipocyte differentiation in mice lacking the C/EBPβ and/or C/EBPδ gene. The EMBO Journal. 16, 7432–7443.

32. Wu, Z., Xie, Y., Bucher, N. L., and Farmer, S. R. (1995) Conditional ectopic expression of C/EBP beta in NIH-3T3 cells induces PPAR gamma and stimulates adipogenesis. Genes & Development. 9, 2350–2363.

33. Tang, Q.-Q., Otto, T. C., and Lane, M. D. (2003) CCAAT/enhancer-binding protein β is required for mitotic clonal expansion during adipogenesis. Proceedings of the National Academy of Sciences. 100, 850–855.

34. Wu, Z., Bucher, N. L., and Farmer, S. R. (1996) Induction of peroxisome proliferator-activated receptor gamma during the conversion of 3T3 fibroblasts into adipocytes is mediated by C/EBPbeta, C/EBPdelta, and glucocorticoids. Molecular and Cellular Biology. 16, 4128–4136.

35. Matsuno, F., Chowdhury, S., Gotoh, T., Iwase, K., Matsuzaki, H., Takatsuki, K., Mori, M., and Takiguchi, M. (1996) Induction of the C/EBP Gene by Dexamethasone and Glucagon in Primary-Cultured Rat Hepatocytes. Journal of Biochemistry. 119, 524–532.

36. Tönjes, R. R., Xanthopoulos, K. G., Darnell, J. E., and Paul, D. (1992) Transcriptional control in hepatocytes of normal and c14CoS albino deletion mice. The EMBO Journal. 11, 127–133.

37. Liu, N., Wei, K., Xun, Y., Yang, X., Gan, S., Xiao, H., Xiao, Y., Yan, F., Xie, G., Wang, T., Yang, Y., Zhang, J., Hu, X., and Xiang, S. (2015) Transcription factor cyclic adenosine monophosphate responsive element binding protein negatively regulates tumor necrosis factor alpha-induced protein 1 expression. Molecular Medicine Reports. 12, 7763–7769.

38. Wang, G., Cheng, Z., Liu, F., Zhang, H., Li, J., and Li, F. (2015) CREB is a key negative regulator of carbonic anhydrase IX (CA9) in gastric cancer. Cellular Signalling. 27, 1369–1379.

39. Lamph, W. W., Dwarki, V. J., Ofir, R., Montminy, M., and Verma, I. M. (1990) Negative and positive regulation by transcription factor cAMP response element-binding protein is modulated by phosphorylation. Proceedings of the National Academy of Sciences. 87, 4320–4324.

40. Chen, W.-K., Kuo, W.-W., Hsieh, D., Chang, H.-N., Pai, P.-Y., Lin, K.-H., Pan, L.-F., Ho, T.-J., Viswanadha, V., and Huang, C.-Y. (2015) CREB Negatively Regulates IGF2R Gene Expression and Downstream Pathways to Inhibit Hypoxia-Induced H9c2 Cardiomyoblast Cell Death. International Journal of Molecular Sciences. 16, 27921–27930.

41. Johannessen, M., Delghandi, M. P., and Moens, U. (2004) What turns CREB on? Cellular Signalling. 16, 1211–1227.

42. Naqvi, S., Martin, K. J., and Arthur, J. S. C. (2014) CREB phosphorylation at Ser133 regulates transcription via distinct mechanisms downstream of cAMP and MAPK signalling. Biochemical Journal. 458, 469–479.

43. Reusch, J. E. B., Colton, L. A., and Klemm, D. J. (2000) CREB Activation Induces Adipogenesis in 3T3-L1 Cells. Molecular and Cellular Biology. 20, 1008–1020.

44. Chrestensen, C. A., Shuman, J. K., Eschenroeder, A., Worthington, M., Gram, H., and Sturgill, T. W. (2007) MNK1 and MNK2 Regulation in HER2-overexpressing Breast Cancer Lines. Journal of Biological Chemistry. 282, 4243–4252.

45. Bell, J. B., Eckerdt, F. D., Alley, K., Magnusson, L. P., Hussain, H., Bi, Y., Arslan, A. D., Clymer, J., Alvarez, A. A., Goldman, S., Cheng, S.-Y., Nakano, I., Horbinski, C., Davuluri, R. V., James, C. D., and Platanias, L. C. (2016) MNK Inhibition Disrupts Mesenchymal Glioma Stem Cells and Prolongs Survival in a Mouse Model of Glioblastoma. Molecular Cancer Research. 14, 984–993.

46. Guo, Z., Peng, G., Li, E., Xi, S., Zhang, Y., Li, Y., Lin, X., Li, G., Wu, Q., and He, J. (2017) MAP kinase-interacting serine/threonine kinase 2 promotes proliferation, metastasis, and predicts poor prognosis in non-small cell lung cancer. Scientific Reports. 10.1038/s41598-017-10397-9

47. Wölfl, S., Martinez, C., and Majzoub, J. A. (1999) Inducible Binding of Cyclic Adenosine 3′,5′-Monophosphate (cAMP)-Responsive Element Binding Protein (CREB) to a cAMP-Responsive Promoter in Vivo. Molecular Endocrinology. 13, 659–669.

48. Dey, A., Nebert, D. W., and Ozato, K. (1991) The AP-1 Site and the cAMP- and Serum Response Elements of the c-fos Gene Are Constitutively Occupied In Vivo. DNA and Cell Biology. 10, 537–544.

49. Sibinga, N. E. S., Wang, H., Perrella, M. A., Endege, W. O., Patterson, C., Yoshizumi, M., Haber, E., and Lee, M.-E. (1999) Interferon-γ-mediated Inhibition of Cyclin A Gene Transcription Is Independent of Individual cis-Acting Elements in the Cyclin A Promoter. Journal of Biological Chemistry. 274, 12139–12146.

50. Tommasi, S., and Pfeifer, G. P. (1999) In Vivo Structure of Two Divergent Promoters at the Human PCNA Locus. Journal of Biological Chemistry. 274, 27829–27838.

51. Nichols, M., Weih, F., Schmid, W., DeVack, C., Kowenz-Leutz, E., Luckow, B., Boshart, M., and Schütz, G. (1992) Phosphorylation of CREB affects its binding to high and low affinity sites: implications for cAMP induced gene transcription. The EMBO Journal. 11, 3337–3346.

52. Xie, J., de Souza Alves, V., von der Haar, T., O’Keefe, L., Lenchine, R. V., Jensen, K. B., Liu, R., Coldwell, M. J., Wang, X., and Proud, C. G. (2019) Regulation of the Elongation Phase of Protein Synthesis Enhances Translation Accuracy and Modulates Lifespan. Current Biology. 29, 737–749.e5.

