## Supplementary material for "Identification of DNA response elements regulating expression of CCAAT/enhancer-binding protein (C/EBP) β and δ during early adipogenesis": Tables

**Table 1.** Primers used for RT-qPCR gene expression analyses.

| Gene | F Primer (5’-3’) | R Primer (5’-3’) |
|---|---|---|
| ***Acaca*** | TGCCTCTGAGAACCCGAAAC | GCCAATCCACTCGAAGACCA |
| ***B2m*** | CTGCTACGTAACACAGTTCCACCC | CATGATGCTTGATCACATGTCTCG |
| ***Cebpa*** | AAACAACGCAACGTGGAGAC | AGTTCACGGCTCAGCTGTTC |
| ***Cebpb*** | CTGCGGGGTTGTTGATGT | ATGCTCGAAACGGAAAAGGT |
| ***Cebpd*** | GCAGCCCCAAAAGCCAGTAAT | GATCAGGGAAGGGGTTGGAA |
| ***Hprt*** | ACATTGTGGCCCTCTGTGTG | TTATGTCCCCCGTTGACTGA |
| ***Mknk1*** | GATTCCTCTGAGACTCCAAGTTAA | ACGCTTCTTCTTCCTCCTCTT |
| ***Mknk2*** | CCAGTGCCAGGGACATAGG | GCCACGCATCTTCTCAAACA |
| ***Nono*** | TGCTCCTGTGCCACCTGGTACTC | CCGGAGCTGGACGGTTGAATGC |
| ***Pparg*** | TCCGTGATGGAAGACCACTCGCAT | CAGCAACCATTGGGTCAGCTCTTG |

**Table 2.** Primary antibodies used for detection of proteins by Western blot. Abbreviations: M: mouse, R: Rabbit.

| Target | Dilution | Species | Catalog. | Supplier |
|---|---|---|---|---|
| **eIF4E** | 1:1000 | R | CST-9742 | Cell Signaling |
| **P-eIF4E (Ser209)** | 1:1000 | R | PA-44528G | ThermoFisher |
| **rpS6** | 1:1000 | M | sc-74459 | Santa-Cruz |
| **P-rpS6 (Ser240/244)** | 1:1000 | R | CST-2215 | Cell Signaling |
| **ERK** | 1:1000 | R | CST-9102 | Cell Signaling |
| **P-ERK (Thr202/Tyr204)** | 1:1000 | R | CST-4370 | Cell Signaling |
| **PKB** | 1:1000 | R | CST-4685 | Cell Signaling |
| **P-PKB (Ser473)** | 1:500 | R | CST-9271 | Cell Signaling |
| **4EBP1** | 1:1000 | R | CST-9644 | Cell Signaling |
| **P-4EBP1 (Ser65)** | 1:500 | R | CST-9451 | Cell Signaling |
| **C/EBPα** | 1:500 | R | CST-2295 | Cell Signaling |
| **C/EBPβ** | 1:500 | R | sc-150 | Santa-Cruz |
| **C/EBPδ** | 1:500 | R | CST-2318 | Cell Signaling |
| **PPARγ** | 1:1000 | R | CST-2443 | Cell Signaling |
| **FABP4** | 1:1000 | R | CST-3544 | Cell Signaling |
| **MNK1** | 1:1000 | R | CST-2195 | Cell Signaling |
| **CREB** | 1:500 | R | CST-9197 | Cell Signaling |
| **P-CREB (Ser133)** | 1:500 | R | CST-9198 | Cell Signaling |
| **ACTIN** | 1:5000 | M | A2228 | Sigma-Aldrich |
| **GAPDH** | 1:1000 | M | G8795 | Sigma-Aldrich |

**Table 3.** Antibodies used for ChIP.

| **Target** | **Dilution** | **Species** | **Catalog.** | **Supplier** |
|---|---|---|---|---|
| **C/EBPα** | 1:100 | R | CST-2295 | Cell Signaling |
| **C/EBPβ** | 1:100 | R | sc-150 | Santa-Cruz |
| **P-CREB** | 1:100 | R | CST-9198 | Cell Signaling |
| **GR** | 1:100 | R | CST-12041 | Cell Signaling |
| **IgG** | 1:1000 | R | CST-2729 | Cell Signaling |
| **PPARγ** | 1:100 | R | CST-2443 | Cell Signaling |

**Table 4.** Primers used for enrichment of protein-bound DNA regions. Gene promoter, type of response element and the distance from transcription start site are indicated. Regions where no specific enrichment was expected are indicated as NEG (negative).

| Protein | Region | F Primer (5'-3') | R Primer (5'-3') |
|---|---|---|---|
| CREB | *Cebpb* CRE -60/106 | CCCCGCGTTCATGCAC | CCACTTCCATGGGTCTAAAGGC |
| | *Cebpb* CRE -2343/2398 | CATGGCCTATTGAGCAAAGAACC | ACCTCATCACAGAGCTTGGC |
| | *Cebpd* CRE -41 | AAACCGCACAAACAGGAAGGAG | ACCGCCGCCTTTTCTAGC |
| | *Mknk1* CRE -60 | AGTGAACTGGCCTTGCTTCTC | GTCGAGAACGCGGAAGAGG |
| | *Mknk2* CRE -372 | AAGGGGAGATTCCGAGGGAAG | AGCCTCTCCACACAGTCCTC |
| GR | *Cebpb* GRE +20 | GCGCCGCCTTATAAACCTC | CAGGCGGTGCATGAACG |
| | *Cebpb* GRE -1161 | GCTAGCGTCTTACCCTTTCCC | CAACCTTCGGTGTTATCTGCTGAG |
| | *Cebpd* GRE -98 | TCCGCCTTTGCTATGTCTGAAG | ACTCCTTGCCTTCCCTCCTTC |
| | *Mknk1* GRE -142 | CTGCCCTCAGGTTTACAAGAACTC | AGAAGCAAGGCCAGTTCACTG |
| | *Mknk2* GRE -86 | ACCAGTCTCCGCCTTTCTCAG | TAAGCTCCGCCCCTTAAACG |
| | *Mknk2* GRE -1883 | TCTGACAGCCAAGTACGTCTTC | TGGTGCCTATAGGGTGAACATC |
| | *Mknk2* GRE -3270 | GTCAGTGTCCTATGCTTGGGATTG | CTTATGTGGTGTGATCTGGTGAGG |
| C/EBPα | *Mknk2* C/EBPRE -1204/1258 | CAGGGGATAGAACTTCAGCTCAAG | CTCTTACATGTGTCCCAGGAATGG |
| C/EBPβ | *Mknk2* C/EBPRE -264/290 | AACACGGCTGCGCACTTC | GGCTGCAGTCGAGTATCTTTTCAC |
| | *Mknk2* C/EBPRE -1204/1258 | CAGGGGATAGAACTTCAGCTCAAG | CTCTTACATGTGTCCCAGGAATGG |
| PPARγ | *Mknk2* PPARE -1975 | ATGTTCACCCTATAGGCACCAG | TGGGAGCCTAGCTTGTAAACAG |
| | *Mknk2* PPARE -3348 | CCATAGAGGAAGAACTGAGACAGC | GGCAGAGCATGTGTCTATTGTACC |
| | *Mknk2* PPARE -3993 | CCATAGAGGAAGAACTGAGACAGC | GGCAGAGCATGTGTCTATTGTACC |
| NEG | *Cebpb* NEG -12460 | TTCCTCAACCTCCTGTCTTGTCTC | GAACTGTACAGCATTCCTGACCAG |
| | *Cebpd* NEG -1378 | GTTGGCGGGTTTCTGAATACAC | CTGGGTCCACTCACTACGTTTATG |
| | *Mknk1* NEG -6325 | GTGGCTTCACCTTTACACATTACC | TTTGTACCTCTGCCCTCTGCTC |
| | *Mknk2* NEG -4862 | GAGTCAACATGGCAGGCTAAAC | GGAGAAGAGAAATATAGGCTTGGG |
